# Proinsulin dependent interaction between ENPL-1/GRP94 and ASNA-1 in neurons is required to maintain insulin secretion in *C. elegans*

**DOI:** 10.1101/2022.06.15.496240

**Authors:** Agnieszka Podraza-Farhanieh, Dorota Raj, Gautam Kao, Peter Naredi

## Abstract

Maturation of insulin is crucial for insulin secretion and function. ENPL-1/GRP94/HSP90B1 plays an important role in this process. ASNA-1/TRC40/GET3 and ENPL-1/GRP94 are conserved insulin secretion regulators in *Caenorhabditis elegans* and mammals and mouse mutants display type 2 diabetes. ENPL-1 and GRP94 bind proinsulin and regulate proinsulin levels in *C. elegans* and cultured cells. Here we found that ASNA-1 and ENPL-1 co-operated to regulate insulin secretion in worms via a physical interaction that required pro-DAF-28/insulin but occurred independently of the insulin binding site of ENPL-1. ASNA-1 acted in neurons to promote DAF-28/insulin secretion. The interaction occurred in insulin expressing neurons and was sensitive to changes in pro-DAF-28 levels. The chaperone form of ASNA-1 is likely bound to ENPL-1. Loss of *asna-1* disrupted Golgi trafficking pathways. ASNA-1 localization was affected in *enpl-1* mutants and ENPL-1 overexpression partially bypassed ASNA-1 requirement. Taken together, we find a functional interaction between ENPL-1 and ASNA-1 which is necessary to maintain proper insulin secretion in *C. elegans* and provides insights about how their loss might produce diabetes in mammals.

## Introduction

ENPL-1 is a *C. elegans* homolog of the endoplasmic reticulum (ER) chaperone - GRP94/GP96/HSP90B1 which has functions both in ER and non-ER compartments ^1–3^. It positively regulates insulin secretion at the level of DAF-28/insulin maturation ^3^. ENPL-1 is likely required to maintain proper homeostasis in the ER since ER stress markers are upregulated in *enpl-1* mutants ^4^. Mammalian GRP94 also helps to maintain proper quality control in the ER of unstressed cells and during the ER-associated degradation ^5,6^. The role of GRP94 and its homologs are essential in organismal development since the deletion of the *Drosophila* gp93 gene led to growth defects ^7^, knockdown in mice caused embryonic lethality ^8^, led to impaired glucose tolerance ^9^, type 2 diabetes ^10^ and defects in the trafficking of the HER2 oncogene ^2^.

The client set of GRP94 is relatively small and restricted to secreted and membrane proteins such as Toll-like receptors ^11,12^, integrins ^13^, insulin-like growth factors ^14,15^ and insulin ^10^. In spite of the variety of roles mediated by GRP94, very few of its co-chaperones and other proteins that modulate its function have been identified, in contrast to its cytoplasmic paralog HSP90. The first identified co-chaperone that is required for GRP94 function is an ER lumen chaperone CNPY3 ^16^. GRP94 interacts with CNPY3 to properly fold TLR proteins, which has functions in innate immunity against microbial infections ^17,18^. Interestingly, the role of CNPY3 was suggested to support the loading of TLR proteins onto GRP94 indicating for the first time that an ER luminal chaperone requires help from other chaperones to fulfil its function. The other well-described chaperone of GRP94 is BiP ^19–22^. GRP94 requires BiP for the accelerating the closure of its open state and trapping the proIGF2 client protein. The slow ATPase activity ^23^ and the slow closure of GRP94 ^22^ suggest that it requires other chaperones to assist in its conformational changes which are required for binding its client proteins. Clients bind via the defined client binding CBD while the regions of GRP94 required for co-chaperone association are unknown.

ASNA-1/TRC40/GET3 is a conserved protein whose function has been mainly associated with transporting tail-anchored proteins (TAPs) to the ER ^24–26^. Work in yeast and worms shows that the ASNA-1/GET3 is found in two redox-sensitive states and that both redox states have distinct functions and structures ^27,28^. Reduced ASNA-1 has a role in inserting TAPs into the ER membrane, whereas the oxidized ASNA-1 is general chaperone with roles in protecting cells from oxidative damage and aggregated proteins ^27,29^. Mutations in human *Asna1/TRC40* have been associated with diseases such as epilepsy and heart development ^30,31^. Furthermore, loss of *asna-1* causes insulin secretion defects in *C. elegans* ^32^ and loss in pancreatic beta cells in mice leads to type 2 diabetes ^33,34^. Although some phenotypes of *asna-1* mutants are associated with the defective TAP insertion, it remains unknown whether the defect in insulin secretion is a consequence of miss-inserted TAPs via its reduced dimeric form or because of loss of functions associated with the oxidized tetrameric form.

Here we show that ENPL-1 and ASNA-1 work together to mediate insulin secretion. The interaction requires pro form of DAF-28/insulin and it is more likely to take place when ASNA-1 is present in the oxidized state. We provide evidence that ASNA-1 binds to ENPL-1 independently of its client binding domain and that increased proinsulin levels promote higher levels of interaction between ASNA-1 and ENPL-1. We find that although both proteins are present in most tissues, most of the interaction between the two proteins occurs in neurons, and specifically in neurons that express the DAF-28/insulin. We also find that overexpression of ENPL-1 can partially bypass the strict block of DAF-28/insulin secretion in *asna-1* mutants. Since ENPL-1 is important for proinsulin binding we show that the interaction of ENPL-1 and ASNA-1 is necessary to maintain proper insulin secretion in *C. elegans*.

## Results

### ENPL-1 and ASNA-1 interact *in vivo* in intact *C. elegans*

In mammals knockdown of the homologs of both ASNA-1/TRC40 and ENPL-1/GRP94 genes result in diabetes ^9,33^. In *C.elegans* both ASNA-1 and ENPL-1 positively promote insulin secretion ^3,32^ and *enpl-1* was identified in a screen for RNAi clones that produced an *asna-1(-)* like phenotype ^35^. As a first step to investigate a possible interaction between the two proteins, we asked if the levels of *asna-1* and *enpl-1* are influenced by the loss of each other. Western blotting and qRT-PCR analysis of *asna-1* levels in *enpl-1(ok1964)* mutants indicated that levels of *asna-1* are unchanged compared to wild-type (Fig. S1A,B). On the other hand, qRT-PCR analysis of *enpl-1* in *asna-1(ok938*) mutants showed that levels of *enpl-1* are significant upregulated (Fig. S1C). Consistently, increased levels of the mouse homolog GRP94, had been previously reported in ASNA1 knockdown mice ^33^. We next asked whether the two proteins might interact to promote insulin secretion function by performing co-immunoprecipitation followed by western blot analysis in strains expressing multi-copy transgene of ASNA-1::GFP and a single copy of 3xFlag::ENPL-1. Both tagged proteins were expressed under their native promoters ^3,32^. This analysis revealed that the two proteins can physically bind (Fig.1A). To account for the possibility of binding between those two proteins after preparation of lysates, we expressed ASNA-1::GFP under control of a pan-neuronal promoter and 3xFlag::ENPL-1 under control of a body wall muscle promoter and performed the co-immunoprecipitation followed by western blot analysis. The results obtained from this experiment indicated that there is no post-lysis interaction since the two proteins did not co-immunoprecipitate when expressed in separate tissues (Fig. S2).

**Figure 1.**
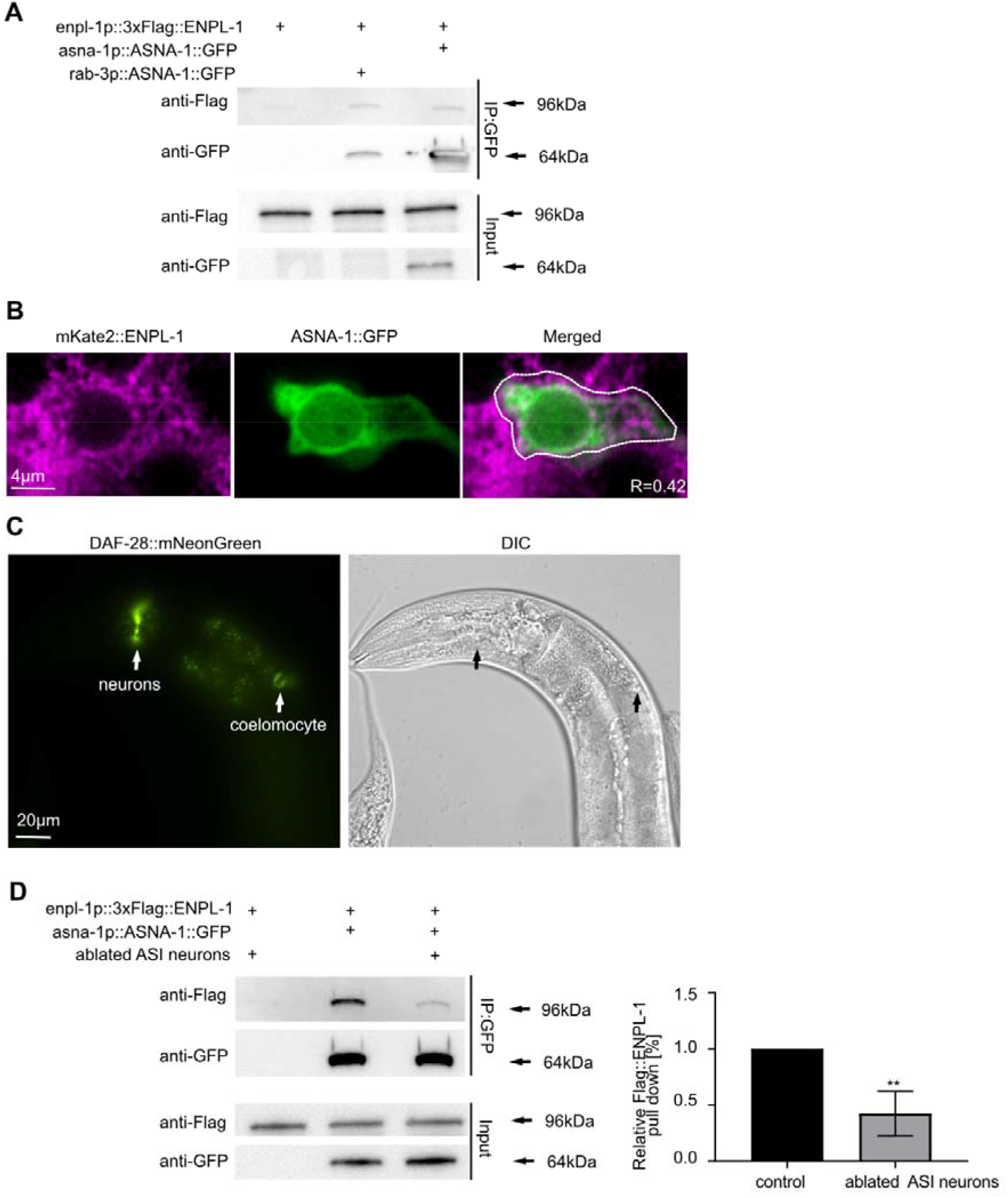
ENPL-1 and ASNA-1 interact in *C. elegans* in insulin expressing neurons. (A) Co-immunoprecipitation experiments with anti-GFP affinity beads from lysates of adult animals expressing 3xFlag::ENPL-1, co-expressing 3xFlag::ENPL-1 + rab3p::ASNA-1::GFP, and 3xFlag::ENPL-1 + asna-1p::ASNA-1::GFP. Inputs from every strain used in the analysis were used as a loading control. (B) Representative confocal images of adult worms expressing ENPL-1::mKate2 and ASNA-1::GFP (n≥12). Scale bar: 4µm. The graph represents Pearson’s correlation coefficient to quantify the localization between ASNA-1::GFP and ENPL-1::mKate2 in neurons. (C) Representative fluorescence microscopy and differential interference contrast (DIC) microscopy images of adult worms expressing DAF-28::mNeonGreen. White arrows indicate locations of expression and secretion of DAF-28::mNeonGreen. Scale bar: 20 µm. (D) Co-immunoprecipitation experiments with anti-GFP affinity beads from lysates of adult animals expressing 3xFlag::ENPL-1, co-expressing 3xFlag::ENPL-1 + ASNA-1::GFP with and without *oyIs84* followed by western blot analysis. Inputs from every strain used in the analysis were used as a loading control. The experiment was performed in triplicate. Quantification shows relative levels of 3XFlag::ENPL-1 immunoprecipitation in the indicated strains. Statistical significance was determined using the two-tailed t-test (**P<0.01). Bars represent mean ± SD.

### ENPL-1 and ASNA-1 interact in the DAF-28/insulin expressing ASI neuron

Since both ENPL-1 and ASNA-1 are required for DAF-28/insulin secretion, we wished to know whether the binding between these two proteins occurred in DAF-28 expressing cells. The DAF-28 protein is found only in two neurons; ASI and ASJ ^32^. To further investigate the localization of a possible ASNA-1 and ENPL-1 interaction, we expressed ASNA-1::GFP under the *prab3* pan-neuronal promoter and performed co-immunoprecipitation in worms co-expressing 3xFlag::ENPL-1 driven by its own promoter. This analysis showed that the same amount of 3xFlag::ENPL-1 was co-precipitated when ASNA-1::GFP was expressed under the neuron specific promoter compared to its own promoter. This indicated that the two proteins interacted in neurons of *C. elegans* and that the bulk of binding occurred in neurons (Fig. 1A). ASNA-1::GFP is expressed in two pairs of neurons of *C. elegans:* ASI and ASJ ^32^ and ENPL-1::mKate2 is widely expressed in neurons ^3^. To further study the neuronal interaction we performed confocal microscopy analysis in worms co-expressing ASNA-1::GFP and ENPL-1::mKate2 and found that both proteins co-localize in neurons (Fig. 1B). Pearson’s correlation coefficient indicated 40% of co-localization between the two proteins (Fig. 1B). We previously showed that insulin/DAF-28::GFP expressed from a multi-copy transgene *svIs69,* is detected in ASI and ASJ neurons, the intestine, and secreted into the coelomocytes ^32^. To ask whether this transgene was accurately reporting the expression of DAF-28, we analyzed a strain in which the mNeonGreen protein was inserted into the *daf-28* genomic locus in order to examine insulin expression and secretion in a non-overexpressed manner. Our analysis showed that the expression of DAF-28::mNeonGreen was still found only in two pairs of head neurons (axons and cell bodies) and the protein was secreted into the coelomocytes from the 4^th^ larval stage onwards (Fig. 1C, Fig.S3A,B). No expression was detected in the intestine or other tissues. To ask if the interaction occurred in DAF-28 expressing neurons we examined the consequences of killing the ASI neurons (33) for the ability of ENPL-1 and ASNA-1 to bind to each other. We found that the immunoprecipitation of 3xFlag::ENPL-1 by ASNA-1::GFP was significantly reduced when ASI neurons were killed (Fig. 1D). Taken together, this analysis supplemented by the confocal analysis indicated that ASNA-1 and ENPL-1 interacted in *C. elegans* and that the interaction took place in ASI neurons which are also a site of DAF-28/insulin expression. We concluded that although both ENPL-1 and ASNA-1 were widely expressed, their interaction might be happening to a large extent only in DAF-28/insulin expressing cells.

### ASNA-1 function is required in neurons to regulate insulin secretion

Having determined that ENPL-1 and ASNA-1 interact in DAF-28 expressing neurons, we next wanted to ask whether ASNA-1 function was required in neurons to promote insulin secretion. For this we used a strain in which we knocked down ASNA-1 protein levels specifically from neurons using the auxin mediated protein degradation system ^36,37^. To do this we used an *asna-1* allele in which mNeonGreen and AID (auxin-induced degron) were inserted at the C-terminus. We analyzed worms expressing a pan-neuronal nuclear tagRFP along with ASNA-1::mNeonGreen::AID and found that ASNA-1::mNeonGreen was expressed in many neurons (Fig. 2A). To deplete ASNA-1::mNeonGreen::AID from neurons we crossed in the *reSi7* transgene that restricts auxin mediated depletion of AID tagged proteins only to neurons ^36,37^. We obtained an efficient auxin mediated knockdown of ASNA-1::mNeonGreen from neurons since we did not observe any ASNA-1::mNeonGreen positive neurons in the nerve ring (Fig. 2B). Next, to understand if the neuronal ASNA-1 was responsible for regulating insulin secretion we analyzed ASNA-1::mNeonGreen::AID;reSi7;DAF-28::GFP worms by exposing them to auxin. Knockdown of ASNA-1 from neurons significantly decreased DAF-28::GFP/insulin secretion (Fig. 2C). Taken together, this data indicated that neuronal ASNA-1 is required for regulation of DAF-28/insulin secretion in *C. elegans*.

**Figure 2.**
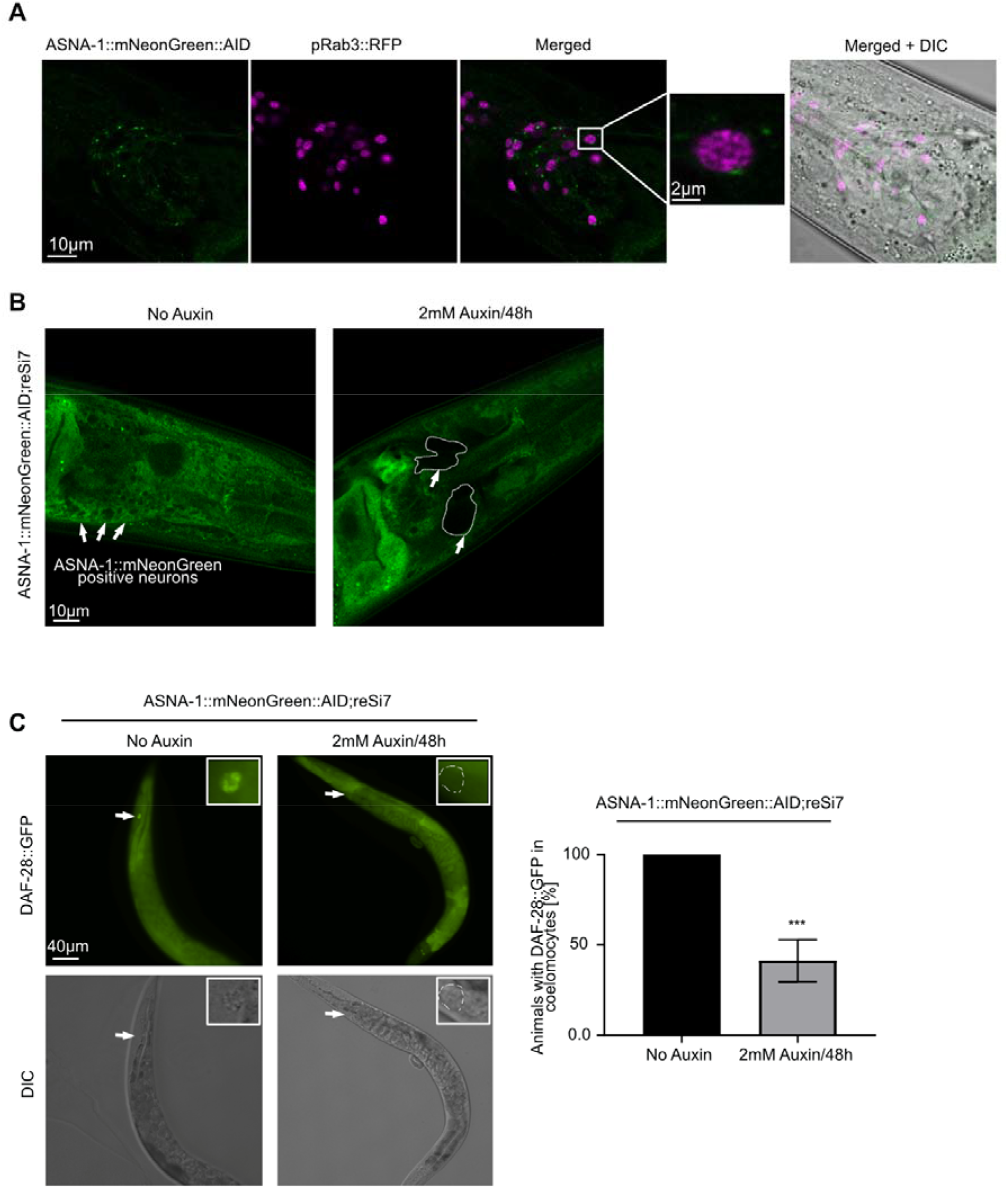
ASNA-1 is required in neurons to regulate insulin secretion. **(A)** Representative confocal images of adult worms expressing pRab3::2xNLS::tagRFP (*otIs364)*, ASNA-1::mNeonGreen and DIC, (n≥10). Scale bar: 10µm. (B) Representative confocal images of adult worms expressing ASNA-1::mNG::AID;reSi7 without auxin or exposed to 2mM auxin for 48h. White arrows indicate ASNA-1::mNeonGreen in neurons, which is absent upon neuronal knockdown (dashed lines). (n≥10). Scale bar: 10 µm. (C) Representative fluorescence and differential interference contrast (DIC) images of worms expressing ASNA-1::mNG::AID;reSi7 with DAF-28::GFP without auxin or exposed to 2mM auxin for 48h. White arrows indicate the localization of coelomocytes in each animal. The dashed line shows DAF-28::GFP-negative coelomocyte. (n≥15). Scale bar: 40 µm. The experiments were performed in triplicate. Quantification shows the percentage of animals with DAF-28::GFP in coelomocytes. Statistical significance was determined using the two-tailed t-test (**P<0.01). Bars represent mean ± SD.

### The interaction between ENPL-1 and ASNA-1 requires pro form of DAF-28

To understand further if the role of ASNA-1 and ENPL-1 interaction was required for maintaining proper insulin secretion, we asked whether insulin was essential for complex formation. We showed previously that ENPL-1 interacted with proinsulin/proDAF-28 in *C. elegans* via its client binding domain. The interaction was essential for the processing of proinsulin to mature insulin since only pro-DAF-28 was detected in *enpl-1* mutants ^3^. Consistent with our findings, it has been shown that murine GRP94 is essential for proinsulin handling and is required for insulin secretion ^10^. Mindful of these findings, we asked if the ASNA-1 and ENPL-1 complex could be affected by the lack of insulin/DAF-28 by performing the co-immunoprecipitation analysis in *daf-28(tm2308)* loss of function mutants ^38^. The interaction between ASNA-1 and ENPL-1 was significantly downregulated in DAF-28/insulin mutants (Fig. 3A). This indicated that the known ENPL-1 client - DAF-28, was also required for the proper formation of ASNA-1 / ENPL-1 complex. To be properly processed and secreted, proinsulin needs to be cleaved in the dense core vesicles by the proprotein convertases 1/3 and 2 ^39^. *C. elegans* insulins also require the activity of proprotein convertases: AEX-5, BLI-4, KPC-1 and EGL-3 ^40–42^. Pro-DAF-28 is cleaved only by KPC-1, whereas INS-4 is cleaved by the EGL-3 ^3,38^. Since the loss of DAF-28 decreases the interaction between ASNA-1 and ENPL-1, we asked whether there would be any effect in the *kpc-1* mutants which accumulate pro-DAF-28 ^3,38^. Co-immunoprecipitation analysis showed that the interaction of ASNA-1::GFP and 3xFlag::ENPL-1 was significantly higher in the *kpc-1* mutants (Fig. 3B). We asked next if another insulin, INS-4, which is processed by EGL-3, similarly affected complex formation by performing co-immunoprecipitation in *ins-4(tm3620)* mutants. In contrast to the findings with *daf-28* mutants, there was no decrease in the immunoprecipitation of ENPL-1 by ASNA-1 pulldown (Fig. 3C). We asked further whether the decreased insulin secretion in the *C. elegans* might affect the strength of binding between ASNA-1 and ENPL-1. For this we used *unc-31(e928)* mutants which display decreased insulin secretion since the UNC-31/CAPS is required for dense-core vesicle release ^43^. In *unc-31* mutants there is accumulation of mature DCVs containing fully processed insulin protein. Co-immunoprecipitation analysis revealed that in *unc-31* mutants, the interaction between ASNA-1 and ENPL-1 was significantly decreased (Fig. 3D). Taken together our analysis showed that DAF-28, and specifically pro-DAF-28 is required for efficient complex formation between ASNA-1 and ENPL-1, and levels of the complex changed with changes in the levels of pro-DAF-28.

**Figure 3.**
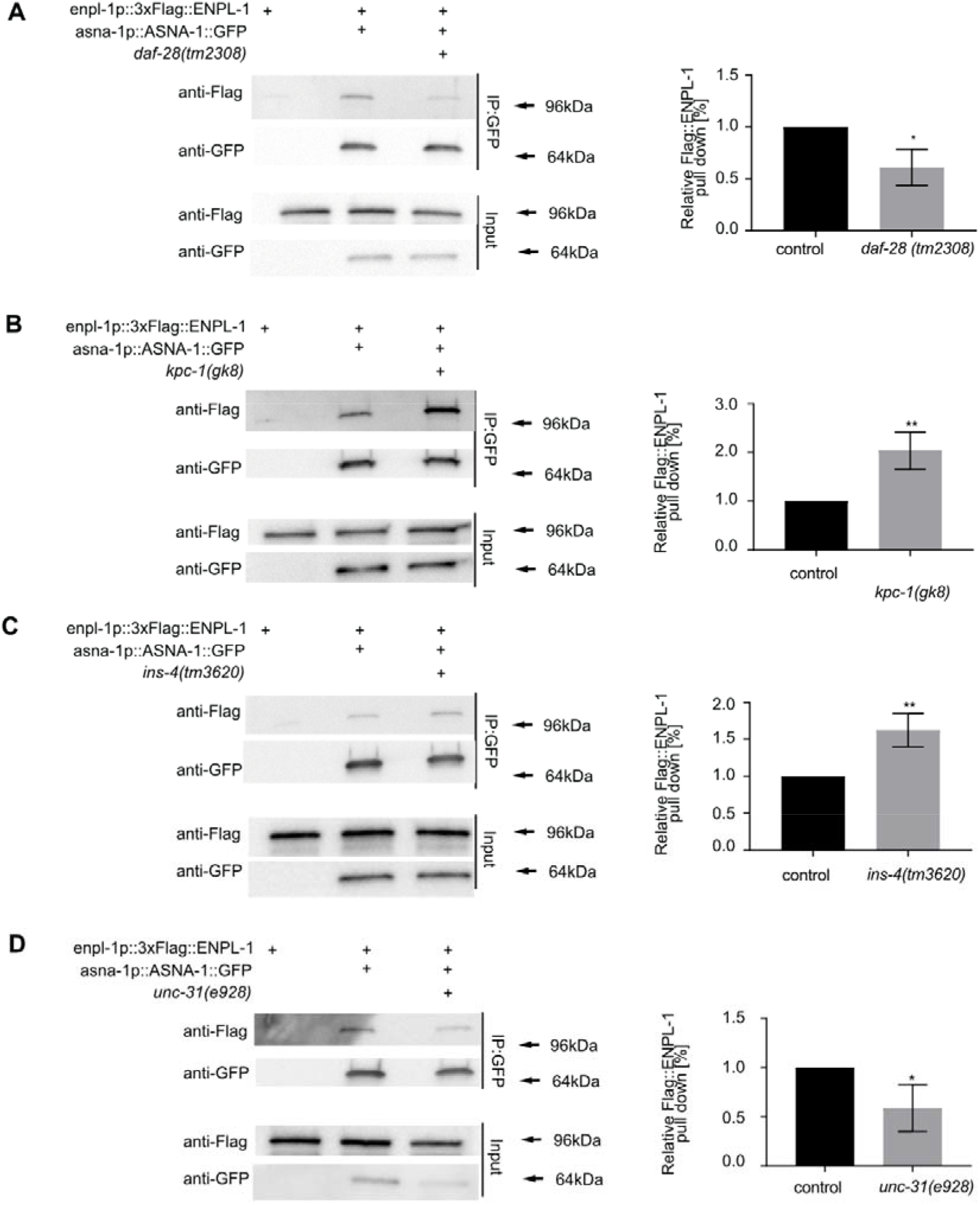
ENPL-1 and ASNA-1 interaction requires proinsulin/pro-DAF-28. Co-immunoprecipitation experiments with anti-GFP affinity beads from lysates of adult animals expressing 3xFlag::ENPL-1 or co-expressing 3xFlag::ENPL-1 + ASNA-1::GFP (A) in *daf-28(tm2308)* mutants, (B) in *kpc-1(gk8)* mutants, (C) in *ins-4(tm3620)* mutants (D) in *unc-31(e928)* mutants, followed by western blot analysis. Inputs from every strain used in the analysis were used as a loading control. The experiments were performed in triplicate. Quantification shows relative levels of 3xFlag::ENPL-1 immunoprecipitation in indicated strains. Statistical significance was determined using the two-tailed t-test (*P<0.05, **P<0.01). Bars represent mean ± SD.

### Increasing levels of oxidized ASNA-1 leads to more interaction with ENPL-1

It has been shown that *C. elegans* ASNA-1 and its yeast homolog, GET3 can work as ATPase-dependent targeting protein. However, under the high oxidative stress condition, the protein undergoes an oxidation dependent conformational change and is converted into an ATPase-independent chaperone ^27,28^. To understand further the character of interaction between ASNA-1 and ENPL-1 we asked if conditions that produce high ROS levels, and convert ASNA-1 to the oxidized form, could influence the interaction with ENPL-1. *sod-2(gk257)* mutants or exposure to H_2_O_2_, are sufficient to increase levels of oxidized ASNA-1::GFP at the expense of the reduced form of the protein ^28^. We performed co-immunoprecipitation in both settings, *sod-2(gk257)* mutants or exposure to H_2_O_2_, and found that in both cases, a significant increase in the interaction between ASNA-1 and ENPL-1(Fig. 4A, B). We concluded that conditions that promote the conversion of ASNA-1 into the oxidized chaperone form promoted increased complex formation, indicating that most likely it is the oxidized form of ASNA-1 which interacts with ENPL-1.

**Figure 4.**
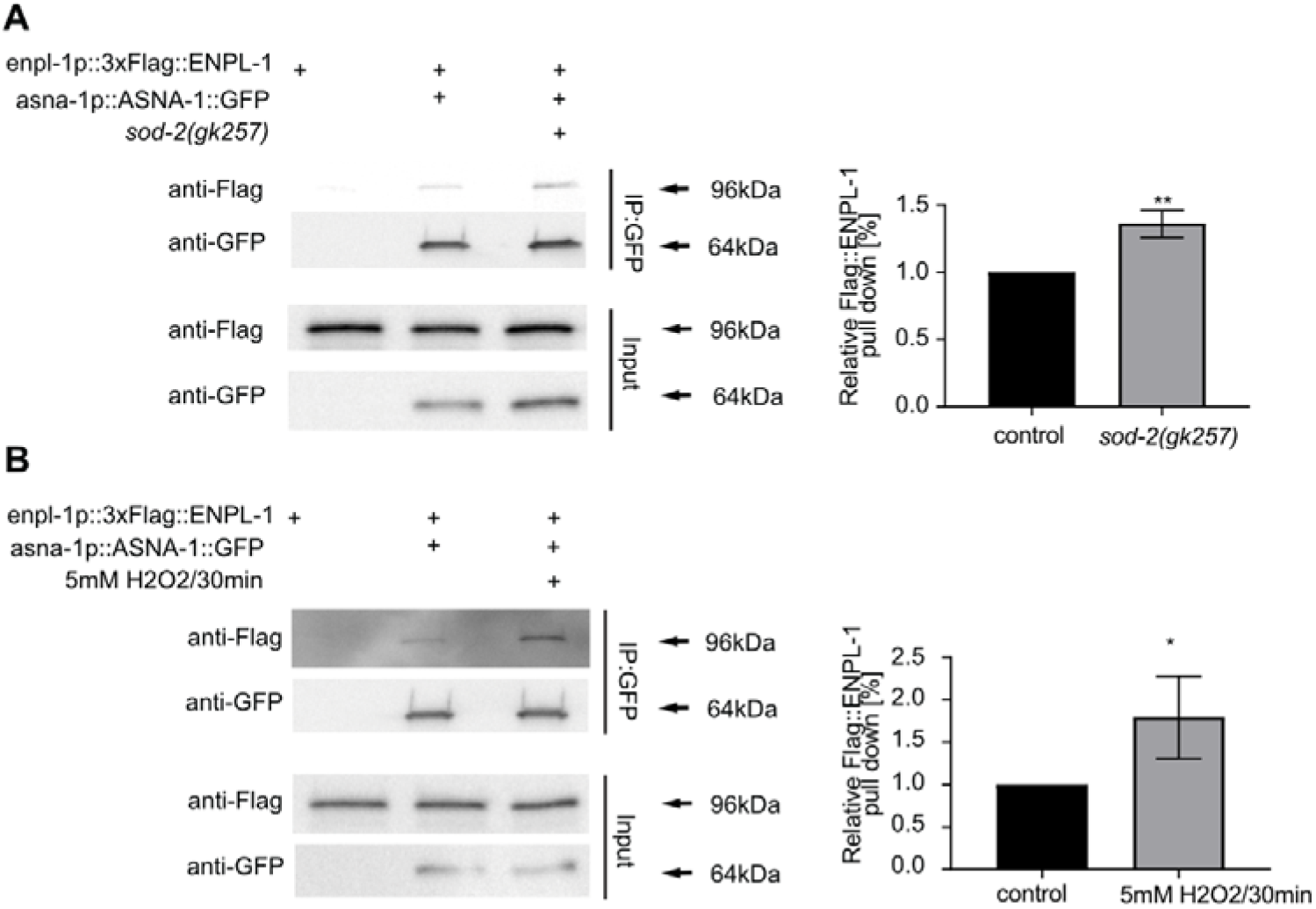
Conditions promoting ASNA-1 oxidation increase ASNA-1/ENPL-1 complex formation. Co-immunoprecipitation experiments with anti-GFP affinity beads from lysates of adult animals expressing 3xFlag::ENPL-1 and co-expressing 3xFlag::ENPL-1 + ASNA-1::GFP (A) in *sod-2(gk257)* mutant, (B) exposed to 5mM H_2_O_2_ for 30min, followed by western blot analysis. Inputs from every strain used in the analysis were used as a loading control. The experiments were performed in triplicate. Quantification shows relative levels of 3xFlag::ENPL-1 immunoprecipitation in the indicated strains. Statistical significance was determined using the two-tailed t-test (*P<0.05, **P<0.01). Bars represent mean ± SD.

### ASNA-1 interacts with ENPL-1 independently of its client binding domain and does not interact with proinsulin

Given our finding that the chaperone form of ASNA-1 was the likely binding partner of ENPL-1, we next asked if ASNA-1 had the characteristics of an ENPL-1 client protein or whether its function was rather based on a non-client role. To address this we used a previously described 3xFlag::ENPL-1 variant, 3xFlag::ENPL-1^ΔCBD^ which has a deletion in the highly conserved client-binding domain (CBD). This deletion was sufficient to prevent its interaction with proinsulin, indicating that the proinsulin is likely a client protein of ENPL-1 ^3^. Upon performing co-immunoprecipitation in worms co-expressing ASNA-1::GFP and 3xFlag::ENPL^ΔCBD^ we found that the CBD from ENPL-1 did not affect the interaction with ASNA-1::GFP, indicating that ASNA-1 was likely not a client protein to ENPL-1, but rather the interaction requires other domains of ENPL-1 (Fig.5A). This finding was consistent with the fact that the client set of GRP94 is restricted to secreted and transmembrane proteins, which ASNA-1 is not. We next asked if ASNA-1::GFP could immunopreciptate proinsulin/pro-DAF-28. To determine this we performed a co-immunoprecipitation experiment from worms expressing ASNA-1::GFP and Ollas::DAF-28::MYC (doubled tagged insulin ^3^) and did not detect any interaction between ASNA-1::GFP and proinsulin (Fig. 5B). We concluded that it is unlikely that there is a direct interaction between the two proteins.

**Figure 5.**
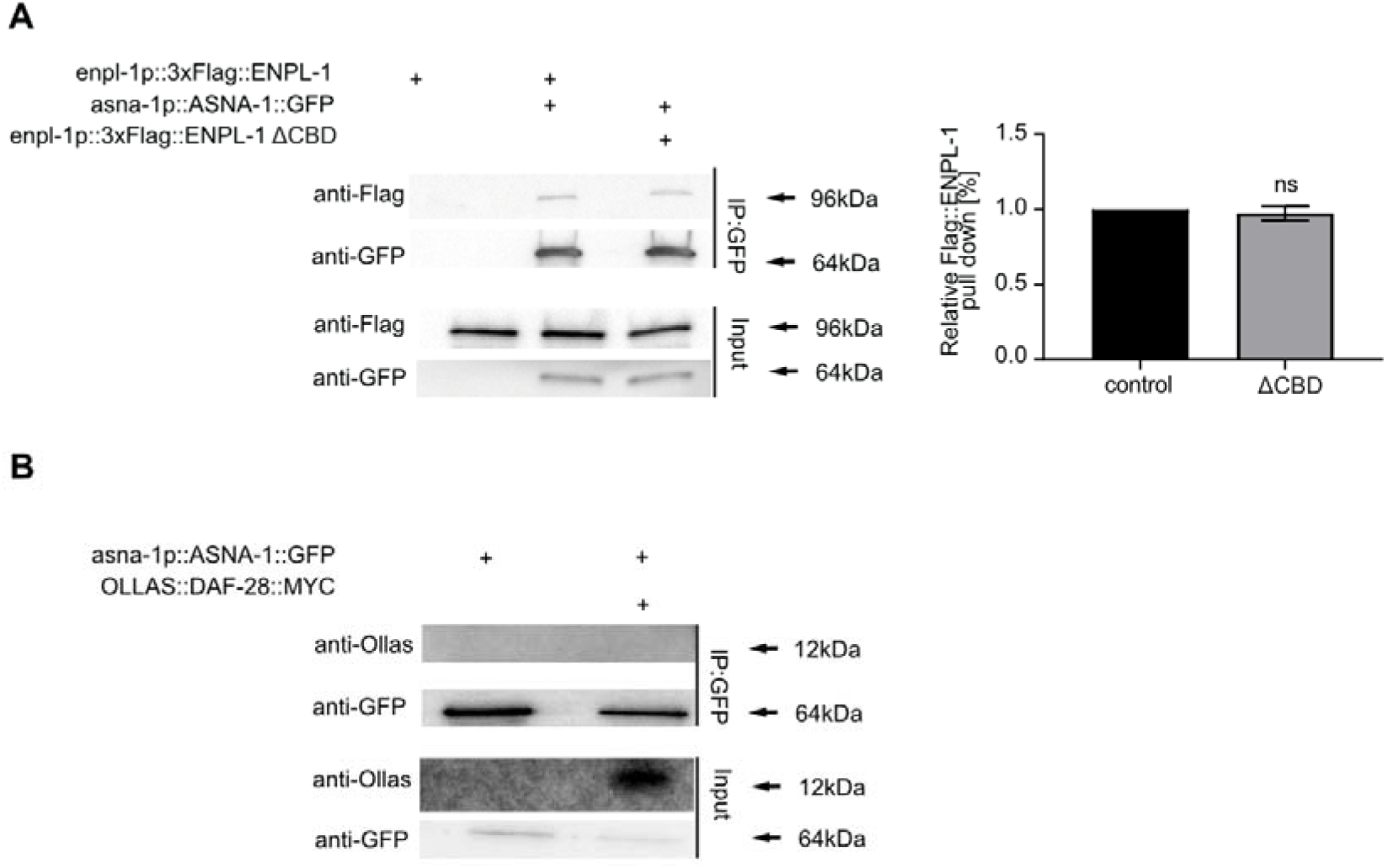
ASNA-1 interacts with ENPL-1 independently of the client binding domain, and does not interact with the pro form of DAF-28. (**A**) Co-immunoprecipitation experiments using anti-GFP affinity beads from lysates of adult animals expressing 3xFlag::ENPL-1, co-expressing 3xFlag::ENPL-1 + ASNA-1::GFP or co-expressing 3xFlag::ENPL-1(ΔCBD) + ASNA-1::GFP followed by western blot analysis. Inputs from every strain used in the analysis were used as a loading control. The experiments were performed in triplicate. Quantification shows relative levels of 3xFlag::ENPL-1 immunoprecipitation in the indicated strains. Statistical significance was determined using the two-tailed t-test. Bars represent mean ± SD. (B) Co-immunoprecipitation experiments with anti-GFP affinity beads from lysates of adult animals expressing ASNA-1::GFP with and without OLLAS::DAF-28::MYC followed by western blot analysis. Inputs from every strain used in the analysis were used as a loading control.

### Overexpression of ENPL-1 bypasses the need for ASNA-1 in DAF-28 secretion

*asna-1* has an essential role in promoting insulin secretion and insulin signaling in *C. elegans* since DAF-28::GFP was not secreted in *asna-1(ok938)* mutants ^32^. Knowing that ASNA-1 and ENPL-1 interact *in vivo* and that the interaction is insulin-dependent, we asked whether overexpression of ENPL-1 could modify the insulin secretion defect of *asna-1(ok938)* mutants. Overexpression of ENPL-1 from 3xFlag::ENPL-1 transgene increases insulin secretion ^3^. We crossed 3xFlag::ENPL-1 into *asna-1(ok938)* mutants and found that overexpression of ENPL-1 partially bypassed the *asna-1* mediated block in insulin secretion. We observed that 26% of animals had improved DAF-28::GFP secretion (Fig. 6A,B). This analysis indicated that there was a compensation of function between these two proteins and that increased levels of ENPL-1 partially suppressed the *asna-1(ok938)* mutant defect.

**Figure 6.**
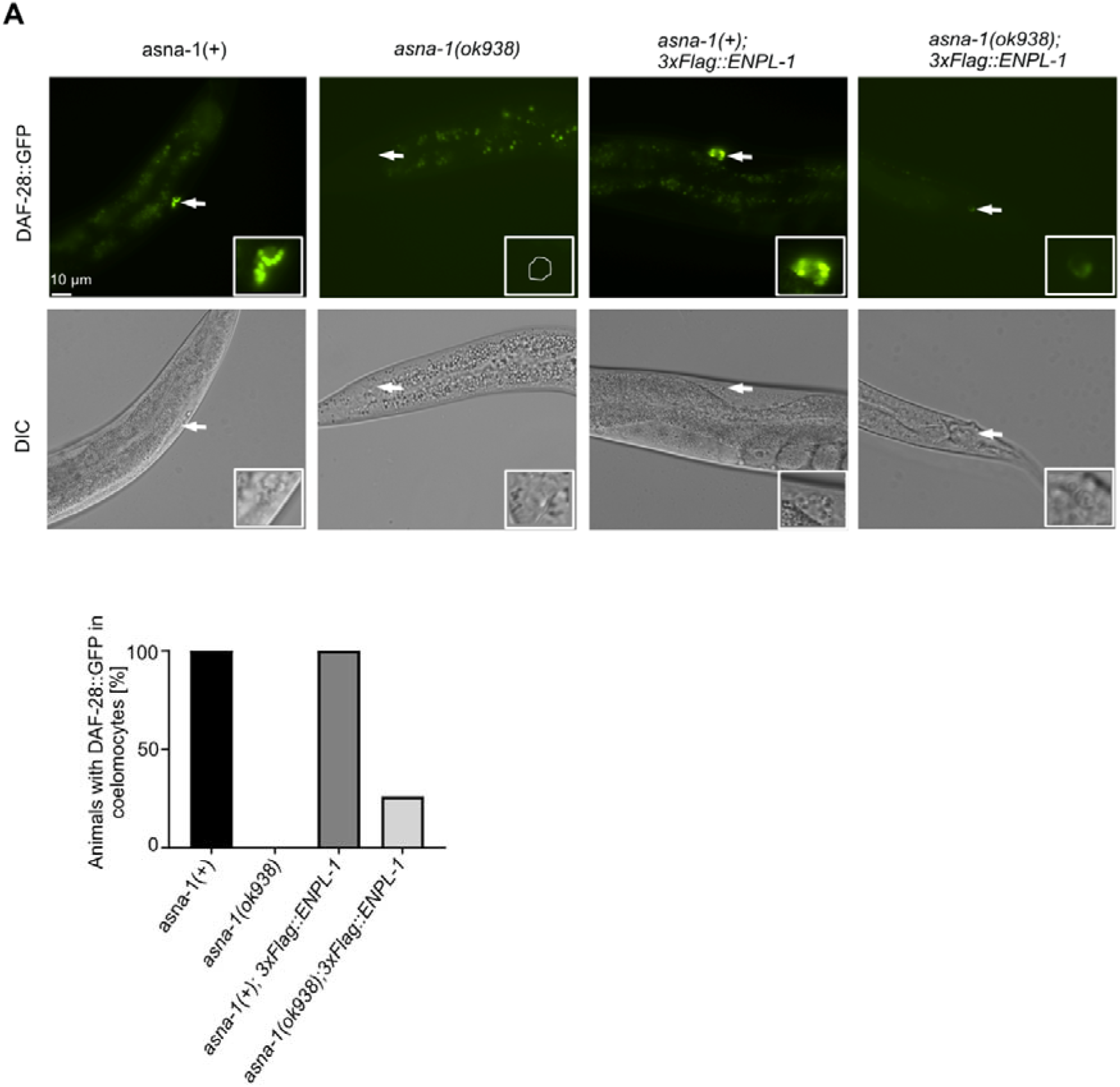
Overexpression of ENPL-1 bypasses the requirement of ASNA-1 for DAF-28::GFP secretion. (A) Representative fluorescence and differential interference contrast (DIC) images of adult *asna-1(+)* and *asna-1(ok938)* animals with and without 3xFlag::ENPL-1. White arrows indicate the coelomocytes accumulating GFP in each animal. The dashed line shows DAF-28::GFP-negative coelomocyte. (n≥10). Scale bar: 10 µm. (B). Bars represent the quantification of the secretion phenotype.

### Loss of *asna-1* perturbs pathways related to ER and Golgi trafficking and transport

To further understand why lack of ASNA-1 causes insulin secretion defects in animals, we performed a quantitative proteomic analysis of *asna-1(ok938)* mutants and wild-type animals to detect proteins and pathways which were the most affected by loss of *asna-1*. Principal component analysis (PCA) indicated distinct expression profiles of wild-type compared to *asna-1(ok938)* mutants (Fig. S4A,B). We detected a total of 4798 proteins among which 1236 were significantly changed (FDR < 0.01) (Fig. S4C). To further understand which pathways were the most affected in the absence of *asna-1*, we performed reactome enrichment analysis and examined the top 20 most upregulated pathways (Fig. S4D). Among them were many pathways related to ER and Golgi transport and trafficking, COP I transport and retrograde transport. Taken together, this data shows that the main role of ASNA-1 is at the level of ER/ Golgi trafficking.

### Loss of *enpl-1* affects the localization of ASNA-1

The proteomic analysis indicated that the role of ASNA-1 was at the level of Golgi function (Fig. S4D). It has been shown as well that the Golgi morphology was affected in the ASNA1knock down mice and led to the formation of small distended membrane stacks ^34^. Knowing this, we asked if the ASNA-1 might be localized to the Golgi. The Golgi markers AMAN-2 and SQV-8 localize Golgi into puncta of a specific morphology in ASI neurons ^44^. Confocal analysis showed that ASNA-1 was widely expressed in embryos, the germline and in neurons. In all these tissues ASNA-1:mNeonGreen is observed in puncta which resemble Golgi bodies (Fig. 7A). The punctate localization was affected by the loss of *enpl-1*, since in the *enpl-1(ok1964)* mutants we observed ASNA-1::mNeonGreen in the more diffused pattern, less localized to the Golgi type puncta (Fig. 7B), even though the expression of *asna-1* gene and protein was unchanged in *enpl-1(ok1964)* mutants (Fig. S1A,B).

**Figure 7.**
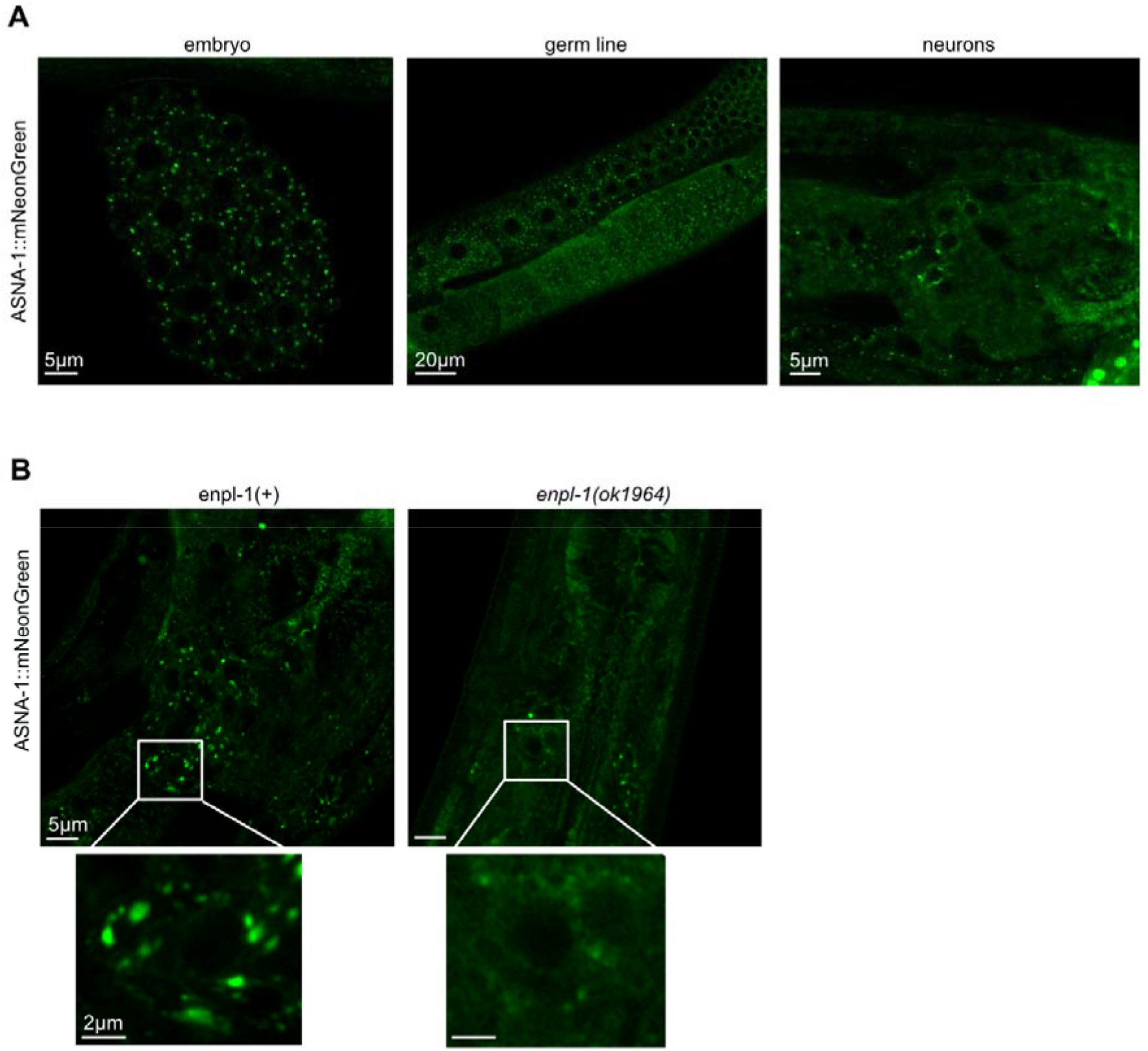
Loss of *enpl-1* causes defects in localization of ASNA-1::mNeonGreen. (A) Representative confocal images of adult worms expressing ASNA-1::mNeonGreen in embryo, germline and neurons. Scale bar: 5µm, 20 µm and 5 µm. (B) Representative confocal images of adult worms expressing ASNA-1::mNeonGreen in *enpl-1(+)* and in *enpl-1(ok1964)* mutants. Scale bar: 5µm, insert 2µm. n≥10 examined for each genotype.

## Discussion

Here we provide the evidence that ENPL-1, the homolog of well-known ER chaperone GRP94/HSP90B1, interacts *in vivo* with ASNA-1. Both ASNA-1 and ENPL-1 are expressed in neurons where the DAF-28/insulin is expressed ^3,32^. Our immunoprecipitation analysis showed that ASNA-1 and ENPL-1 interact in neurons. Most of the interaction occurred in ASI neurons and the interaction required the presence of DAF-28. When there were high levels of pro-DAF-28 in *kpc-1* mutants ^3^, we found increased interaction between ASNA-1 and ENPL-1 indicating that the proinsulin form of DAF-28 likely drives increased binding. Conditions which increased oxidation of ASNA-1 increased the binding between ASNA-1 and ENPL-1 indicating that the oxidized form of ASNA-1 which acts as a chaperone ^27^, binds to ENPL-1. Our study showed that the client binding domain of ENPL-1 which binds proinsulin was not needed for ASNA-1 binding, since the pull-down efficiency was not affected when CBD was mutated. Interestingly, while the ENPL-1 formed a complex with pro-insulin, an ASNA-1/pro-DAF-28 complex was not detected.

Numerous studies have shown that GRP94 is an important ER chaperone, that binds and ensures the folding of essential client proteins such as toll-like receptors, proinsulin or insulin-like growth factor II ^3,10,14,16^. However, it remains unknown how many non-client proteins are needed by GRP94 to maintain its proper function. The ATPase activity of GRP94 is relatively low, indicating that GRP94 requires a co-chaperone to enhance its binding to the designated clients ^23,45^. Recent studies have shown that BiP is a novel interacting partner of GRP94, and that BiP assists with the acceleration of the closure of GRP94 to trap the client protein ^22^. However, there is a lack information on which co-chaperones might act with ENPL-1/GRP94 for proinsulin handling and processing. Our studies in *C. elegans* identified ASNA-1 as new interactor of ENPL-1 and provided evidence consistent with the notion that ASNA-1 might be a co-chaperone of ENPL-1.

Both ASNA1 and GRP94 act to positively regulate insulin secretion in mice ^9,34^. These studies showed that both proteins were essential for pancreas development and function, and specific knockdown in pancreas led to a diabetic phenotype ^9,33,34^. Our studies in *C. elegans* also showed that both proteins, which are well conserved with their mammalian counterparts, are positive regulators of insulin signaling. Notably, human ASNA1 can substitute for worm ASNA-1 for both growth and cisplatin detoxification functions ^32,46^ indicating conservation of function and that results obtained with worm ASNA-1 can be relevant for human biology. Knockdown of either of these two worm genes resulted in growth arrest and a severe defect in insulin secretion ^3,32^. We have previously shown that ENPL-1 binds insulin, and in the absence of *enpl-1*, proinsulin levels are downregulated and only exists in the unprocessed form. This very likely contribute to the defect in insulin secretion. However not much is known at the subcellular level about how ASNA-1 regulates insulin secretion in *C. elegans*. Our analysis here indicated that ASNA-1 expressed specifically in neurons is responsible for regulating insulin secretion in *C. elegans* since the protein knockdown resulted in insulin secretion defect. This data together with our demonstration of the interaction between ASNA-1 and ENPL-1 in insulin specific neurons supports the notion that ASNA-1 is required by ENPL-1 to maintain insulin secretion. We note further that our previous work has shown that ENPL-1 has functions outside the ER compartment ^3^. This provides possible subcellular locations for the interaction between the two proteins. Roles outside the ER are not a feature just of worm ENPL-1. Rather, there is substantial evidence for such non-ER roles for mammalian GRP94/HSP90B1 ^1,2^. Furthermore, it has been also shown that yeast and mammalian homologs of ASNA-1 are required for maintaining proper retrograde transport and retrieval of proteins with the HDEL motif from the Golgi to the ER ^33,47^. We note that ENPL-1 also has a C-terminal HSEL motif, required for its retrieval to the ER ^48^. Our global proteomic analysis reported Golgi and ER trafficking and retrograde transport changes in the *asna-1* mutants as one of the main pathways affected by the loss of ASNA-1.

Our results showed that interaction between ASNA-1 and ENPL-1 is insulin/DAF-28 dependent since in the *daf-28* mutants we observed a significant reduction of the interaction. Our previous studies showed that ENPL-1 interacts with proinsulin, pro-DAF-28 and that in the absence of *enpl-1*, proinsulin cannot be properly cleaved ^3^. This data and additionally the fact that both proteins are essential for insulin secretion indicates that the interaction is related to insulin binding and possibly processing. Although the genome of *C. elegans* encodes for 40 insulin-like proteins ^49^, we have chosen to study two insulins *daf-28* and *ins-4* and their impact on interaction since it has been shown that during maturation, these two insulins are cleaved by two different proprotein convertases, DAF-28 is cleaved by KPC-1, whereas INS-4 is cleaved by EGL-3 ^38^. Additionally, our previous data indicated that, pro-DAF-28 is not cleaved in *kpc-1* mutants, leaving the insulin in the unprocessed form ^3^ which is also consistent with earlier findings ^38^. Interestingly the absence of these two insulins had different impacts on the interaction between ASNA-1 and ENPL-1. Firstly, we observed increased interaction in the *kpc-1(gk8)* mutants, indicating that the proinsulin form of DAF-28 is required for the binding between ASNA-1 and ENPL-1. Secondly, we found that in the absence of *ins-4*, the interaction level increases possibly via an increase in DAF-28 levels. These different results might indicate that the interaction between ASNA-1 and ENPL-1 is directed towards specific insulin – DAF-28, and not all 40 insulins expressed by *C. elegans*. It remains likely that other worm insulins like INS-2 that require KPC-1 for their processing might also require the ENPL-1/ASNA-1 complex for their function ^38^. We do not rule out the possibility that client proteins that do not need KPC-1 for their function might use the ASNA-1/ENPL-1 complex.

ASNA-1 has been mainly described as a protein that promotes the insertion of tail-anchored proteins into the ER ^24–26^. The only known interactors of ASNA-1 or its homologs are proteins that take part in TAP insertion ^50^. However, it has been shown that ASNA-1 and its yeast homolog, Get3, can act as a holdase chaperone that under oxidative stress conditions protect proteins from oxidative stress damage ^27–29,51^. Those changes involve structural rearrangements of the protein including disulfide bond formation, zinc release and oligomerization ^27^. Our previous study showed that *C. elegans* ASNA-1 is normally present both in the oxidized and reduced state in the animal. The oxidized state is favored under the high ROS conditions ^28^. Forcing ASNA-1 to adopt the oxidized state impairs TAP insertion but does not affect insulin secretion indicating that those two functions can be separated and that oxidized ASNA-1 is responsible for the role in insulin secretion ^28^. Our results showed that the interaction between ASNA-1 and ENPL-1 increases under high ROS conditions which lead to oxidation of ASNA-1 and impaired TAP insertion function. This allows us to propose that the oxidized form of ASNA-1 is more likely the binding partner of ENPL-1. More broadly GRP94 has diverse roles that have an impact on human health. The discovery that ASNA-1 is functionally linked to it can be a starting point for genetic or drug based interventions that modulate its activity.

Insulin secretion is central in organism metabolism and for normal life. Much is known of how insulin secretion can be stimulated (e.g. by glucose) but considerably less is known about insulin maturation. Our study links ENPL-1 and ASNA-1 and provides information for how they might cooperate for insulin maturation and secretion and prevent diabetes.

## Materials and Methods

### *C. elegans* genetics and maintenance

Animals were maintained under the standard conditions at 20°C on nematode growth media (NGM) plates. N2 is the wild-type parent for all the strains in the study. The *daf-28::mNeonGreen (PHX3050*) strain was created by SunyBiotech using the CRISPR/Cas9 technique by inserting flexible linker and mNeonGreen before the stop codon of *daf-28*. The multi-copy ASNA-1::GFP(*svIs56*) transgenic animals, the multi-copy DAF-28::GFP(*svIs69*) transgenic animals and *asna-1(ok938)* mutants which were maintained *in trans* to the hT2(*qIs48*) balancer have been previously described in ^32^. The single-copy 3xFlag::ENPL-1(*knuSi222*), the single copy 3xFlag:: ENPL-1(ΔCBD) *knuSi430* with an in-frame deletion of the Client binding domain, the knock-in strains ENPL-1::mKATE2:enpl-1(*PHX698*) and the double-tagged ollas::DAF-28::Myc(*rawEx11*) were previously described ^3^. The ASNA-1::mNeonGreen::AID [*PHX2249*] was created by CRISPR/Cas9 knock-in of two tags mNeonGreen and AID at the C-terminus of the protein of ASNA-1 by Sunybiotech. *enpl-1(ok1964)* was outcrossed 8 times and maintained in trans to the nT1(qIs51) balancer, *sod-2(gk257)*, *daf-28(tm2308)*, *ins-4(tm3620)*, *kpc-1(gk8)*, *unc-31(e928)*, *otIs356* expressing nuclear localized pan neural tag:RFP and *oyIs84* obtained from the *Caenorhabditis* Genetics Center (University of Minnesota).

### Antibodies and western blotting

Wild-type animals or indicated mutants were directly washed using M9 from the plates. Animals were lysed in Next Advance Bullet Blender Homogenizer in buffer containing 10 mM Tris-HCl (pH 7.4), 150 mM NaCl, 5 mM EDTA, 0.5%NP40 (between 80 µl and 200 µl) using 0.2 mm stainless steel beads for 3 min at 4°C, followed by centrifugation at 14,000 rpm (18,400 g) for 20 min at 4°C. Protein estimation was conducted using the BCA assay. *Reducing SDS-PAGE* was performed as described in ^28^. Proteins were separated by SDS-PAGE and blotted onto nitrocellulose membranes. Proteins were detected using the following antibodies: anti:ASNA-1^32^, anti-GFP [3H9, ChromoTek, RRID: AB_10773374, Cat# 3h9-100, LOT# 80626001AB, dilution: 1:1000], anti-GRP94, anti-Flag [M2, Sigma-Aldrich, RRID: AB_262044, Cat# F1804, LOT# SLCD3990, dilution: 1:1000,] anti-tubulin [Sigma-Aldrich, RRID: AB_477579, Cat# T5168, LOT# 00000089494, dilution: 1:5000], anti-OLLAS [L2, Novus Biologicals, RRID: AB_1625980, Cat# NBP1-06713SS, dilution: 1:1000]. The secondary antibodies, used at a dilution of 1:5000, were HRP conjugated: goat anti-rat [GE Healthcare Life Sciences, RRID: AB_772207, Cat# NA935, LOT# 16918042], sheep anti-mouse [GE Healthcare Life Sciences, RRID: AB_772193, Cat# NA9310, LOT# 16921365] and donkey-anti-rabbit [GE Healthcare Life Sciences, RRID: AB_2722659, Cat# NA934, LOT# 9670531]. Supersignal West Femto detection reagent (Thermo Fisher Scientific) was used to generate a signal, which was detected using a LAS1000 machine (Fujifilm).

### Co-immunoprecipitation

Worms were grown on the NGM plates for 4 days at 20°C and lysed as described above. Then 800-1000 µg of total protein lysates were added to 25 μl of anti-GFP magnetic beads (ChromoTek GFP-Trap) and tumbled end-over-end for 1 h, at 4°C. Beads were separated with a magnet and washed 3× for 10 min with 600 μl of wash buffer [10 mM Tris-HCl (pH 7.4), 150 mM NaCl, 5 mM EDTA]. Proteins were eluted by re-suspending the washed beads in 20 μl of 2× loading dye with β-mercaptoethanol, followed by heating for 10 min at 95°C. SDS PAGE was performed as described above.

### Neuropeptide secretion assays

One day adult animals were anaesthetized using 10 mM levamisole, mounted on 2% agarose pads and directly imaged. DAF-28::GFP uptake into coelomocytes of adult worms was measured directly after preparing the slide. Control samples and indicated mutants were measured using the same settings in parallel.

### H_2_O_2_ exposure

Hermaphrodites were grown on the NGM plates for 4 days. After the indicated time animals were washed from the NGM plates and incubated with 5mM H_2_O_2_ for 30 min. Worms were harvested and proceeded for lysate preparation and co-immunoprecipitation.

### Auxin mediated depletion

Water soluble auxin (Naphthaleneacetic Acid (K-NAA)), PhytoTech, LOT#HKA0610009, CAS#15165-79-4 plates were prepared on the day of use by adding indicated concentration of auxin to NGM plates after cooling the agar down to 56°C. The plates containing auxin where kept in the darkness during preparation and experimental procedure. ASNA-1::mNeonGreen::AID;reSi7 animals were grown and staged to obtain l1-l2. Staged animals were put on 2mM auxin plates for 48h and kept in the darkness. After indicated time, fluorescence microscopy and DIC pictures were taken.

### Confocal microscopy

One day old adult animals were anaesthetized using 10 mM levamisole, mounted on 2% agarose pads and directly imaged. The fluorescence signal was analyzed at 488 nm and 561 nm using a confocal laser scanning microscope (LSM880, Carl Zeiss) with C-Apochromat 40x/1.2 water immersion objective lens. Image processing was performed using ZEN Lite (Carl Zeiss) software.

### RNA extraction and quantitative RT-PCR (qPCR)

Worms were grown on the NGM plates for 4 days at 20°C. Worms were synchronized by allowing a mixed-stage worm suspension in M9 buffer to settle for 3 min and collecting the supernatant which contained embryos and L1 larva. These were placed on fresh NGM plates for 48 h to grow until the young adult stage. Worms were re-suspended in 75 µL Nucleozol (Macherey-Nagel). After lysis by three rounds of freeze/thaw (37°C and ethanol/dry ice), the RNA was extracted using the Aurum Total RNA Mini Kit (Bio-Rad). cDNA was synthesized using the iScript cDNA Synthesis Kit (Bio-Rad). qPCR was performed on a StepOnePlus Real-Time PCR System (Applied Biosystems) instrument using KAPA SYBR FAST qPCR Kit (Kapa Biosystems). The comparative Ct method was used to analyze the results and the reference gene used for the analysis was *CDC-42*.

### Proteomic analysis

Staged, adult N2 and *asna-1(ok938)* animals were collected and lysed in 2% SDS in 50mM of Triethylammonium bicarbonate (TEAB) buffer. Total protein estimation was estimated using BCA estimation.

Samples were subjected for Relative Quantitative Mass Spectrometry using TMT performed by The Proteomics Core Facility at Gothenburg University and analyzed by The Bioinformatics Core Facility at Gothenburg University.

In brief:

The samples were homogenized and a reference pool containing equal amounts of all samples was produced representing a mean of the other samples. Equal amounts of proteins from each sample and the reference pool were trypsin digested into peptides. The peptides were subjected to isobaric mass tagging reagent TMT with a unique tag for each sample and the reference. After labeling, samples were combined resulting into one or several 10-plexed sets including the reference pool. The reference pool allows comparison between the different sets. The peptides analyzed by nano-liquid chromatography (LC) on-line coupled to an Orbitrap mass spectrometer (MS). The mass-to-charge (m/z) ratio of the peptides (MS) were determined followed by fragmentation (MS/MS) for peptide sequence information and relative quantification. The analysis was performed in MultiNotch mode where the quantification occurs in MS3 reducing the interference of co-isolated peptides. Relative quantification was performed using Proteome Discoverer or MaxQuant. MS-raw data for each set were merged during the database search for protein identification and relative quantification. Since the tags have an isobaric chemical structure, the peptides labelled with different tags were indistinguishable during chromatographic separations and in MS mode. Each tag contains a characteristic so-called reporter ion with a unique structure which is detectable upon fragmentation. The ratios of these reporter ion intensities in MS/MS spectra were used for quantitation. Only peptides unique for the specific protein are considered for quantitation. Fold change calculation between groups and statistical analysis were performed using Welch’s t-test together with pathway analysis using Ingenity Pathway Analysis to determine the biological relationships, mechanisms, functions, and pathways of relevance for the identified proteins. The low variability and the high sensitivity of the method allow fold-change of >1.2 if p-value is 0.025-0.05 and a fold-change>1.1 if p-value is <0.025 to be considered as significant results.

### Colocalization analysis

Colocalization analysis was performed using Coloc 2 and pixel intensity correlation over space methods of Pearson was used (Fiji Software).

### Statistical analysis

Statistical analysis was performed using Prism 9 software (GraphPad Software). Statistical significance was determined using a two-tailed, unpaired Student’s t-test. P-values <0.05 indicated statistical significance (*P<0.05, **P<0.01, ***P<0.001, ****P<0.0001).

## Acknowledgements

We thank the *Caenorhabditis* Genetic Center (funded by National Institutes of Health Office of Research Infrastructure Programs P40 OD010440) for providing strains, the Centre for Cellular Imaging at the University of Gothenburg and the National Microscopy Infrastructure, NMI (VR-RFI 2016-00968) for providing assistance in microscopy. Bioinformatics and Proteomics Core Facility of Sahlgrenska Academy, the University of Gothenburg for proteomic analysis.

